# Bridging Genomics and Clinical Medicine: RSVrecon Enhances RSV Surveillance with Automated Genotyping and Clinically Important Mutation Reporting

**DOI:** 10.1101/2025.06.03.657184

**Authors:** Lei Li, Haidong Yi, Jessica N. Brazelton, Richard J. Webby, Randall T. Hayden, Gang Wu, Diego R. Hijano

## Abstract

**Background:** Respiratory Syncytial Virus (RSV) causes significant respiratory infections, particularly in young children and elderly adults. Genetic variations in the fusion (F) protein can reduce the efficacy of vaccination and monoclonal antibody treatments, emphasizing the need for genomic surveillance of this virus.

**Motivation:** Current pipelines for RSV genome assembly focus on sequence reconstruction but often lack features for detecting genotypes, clinically relevant mutations, or presenting results in formats that are suitable for clinical researchers.

**Results:** We introduce RSVrecon, an advanced bioinformatics pipeline for comprehensive RSV genome assembly and phylogenetic analysis. RSVrecon processes raw FASTQ files into annotated variant reports and delivers results in multiple formats (CSV, PDF, HTML) tailored to diverse end users. A key innovation of RSVrecon is not only its integrated detection of clinically critical features—including genotype classification and F protein mutation calling, capabilities absent in most analytical pipelines—but also its presentation of these results to clinicians via an integrated, graphical, and user-friendly interface. Its modular design, powered by Nextflow’s modern framework, ensures a scalable and robust workflow, while user-friendly reports enable seamless translation of genomic data into actionable clinical insights. Benchmarking against existing pipelines using clinical datasets revealed that RSVrecon achieves comparable genomic assembly accuracy while excelling in three key dimensions: (1) expanded functional capabilities, (2) intuitive biological interpretation of the results, and (3) superior user experience and accessibility. By seamlessly translating RSV genomic data into clinically meaningful information, RSVrecon empowers research breakthroughs, guides clinical care decisions, and strengthens surveillance systems. With these features, RSVrecon offers an enhanced approach to RSV surveillance and research. The tool is freely available at https://github.com/stjudecab/rsvrecon and https://github.com/stjudecab/RSVreconPy.

**Key Points:**

- RSVrecon enables comprehensive clinical detection for RSV such as identifying genotypes and F protein mutations.
- RSVrecon delivers results through an integrated, graphical, and user-friendly interface for clinicians.
- RSVrecon’s Nextflow-based design ensures robust, scalable, and consistent performance.
- RSVrecon matches assembly accuracy while excelling in functionality, interpretation, and accessibility during benchmarking.

## Introduction

Respiratory infectious diseases are major global pandemic threat (Biggerstaff et al., 2014). Throughout modern history, the most devastating pandemics have been of respiratory origin, including the 1918 influenza pandemic, which infected an estimated 500 million individuals and resulted in 50–100 million fatalities (Morens and Fauci, 2007), the 2009 H1N1 pandemic, which spread to 214 countries within a year and caused an estimated 150,000–200,000 deaths (Dawood et al., 2012), and the COVID-19 pandemic, which led to an estimated 18.2 million deaths in its first two years (2020 - 2021) (Wang et al., 2022). The continuous evolution of influenza is driven by antigenic drift, facilitating seasonal epidemics (Li et al., 2020a, Han et al., 2019, Smith et al., 2004), while antigenic shift underlies the emergence of novel pandemic strains, as observed in the H2N2 (1957), H3N2 (1968), and H1N1 (2009) pandemics (Petrova and Russell, 2018). The COVID-19 pandemic further underscored the pandemic potential of coronaviruses (*Coronaviridae*), whose broad host range and high recombination rates position them alongside influenza viruses as major global health threats (Li et al., 2020b, Goldstein et al., 2022).

Given this demonstrated pandemic potential of respiratory viruses, Respiratory Syncytial Virus (RSV) – though historically understudied – warrants similar attention as an emerging public health challenge. Recent research has begun to reveal its significant disease burden: RSV causes 33 million infections annually and 65,000 deaths in children under five years old, accounting for 20% of global pediatric lower respiratory mortality (Kaler et al., 2023). It also poses a severe risk to older adults, with an 8% mortality rate among hospitalized individuals aged 65 and older (Falsey et al., 2005). Vaccination programs and monoclonal antibodies (mAb) targeting high-risk groupshave been effective in preventing severe RSV (Langedijk et al., 2022, Simões et al., 2021, Zhu et al., 2018, Zhu et al., 2017). However, the efficacy of these interventions is significantly influenced by genetic variations in the virus genome, particularly in antigenic regions of surface proteins. The fusion protein (F protein) is a glycoprotein located on the surface of RSV particles and is critical for the virus’s entry into host cells (Neal et al., 2024, Battles and McLellan, 2019). Several key residues of the F protein have been identified as antibody-binding sites for commonly used mAbs. Substitutions at these residues have been observed to diminish the effectiveness of mAb treatments (Zhu et al., 2018, Zhu et al., 2017, Zhu et al., 2012). Consequently, assembling genomic sequences of RSV from clinical samples and identifying clinically relevant genomic variations are essential for improving clinical treatment and monitoring the evolution of the virus. This approach enables the detection of mutations that may impact therapeutic efficacy and supports the development of more effective interventions.

Sequencing viral samples using next-generation sequencing (NGS) is a relatively mature process and has been extensively studied both experimentally and computationally(Antipov et al., 2022, Chevreux, 2007, Fu et al., 2024, Jansz and Faulkner, 2024, Köndgen et al., 2024, Váradi et al., 2022, Wan et al., 2015, Yamashita et al., 2016, Yang et al., 2012, Zsichla et al., 2024). Numerous computational pipelines have been developed to assemble viral genomes from NGS data. Existing pipelines for RSV genome assembly can be categorized into three main groups based on their core algorithms: de novo-based approaches, reference-based approaches, and hybrid approaches that combine both strategies(Zsichla et al., 2024). In terms of application, there are general genome assembly tools such as CLC Workbench, VirAmp, viralFlye, MIRA, and VICUNA (Wan et al., 2015, Antipov et al., 2022, Chevreux, 2007, Yang et al., 2012, QIAGEN, 2025), alongside RSV-specific methods like NEXT-RSV-PIPE and RSV-GenoScan (Dosbaa et al., 2024, Köndgen et al., 2024). Although these tools deliver accurate sequence assemblies and annotations, they predominantly focus on genomic reconstruction and often fail to address clinically relevant features of RSV such as genotypes and surface protein substitutions. Furthermore, these methods frequently lack features designed for accessibility by clinical researchers and medical professionals. Finally, implementation limitations constrain the usability of existing pipelines, particularly concerning workflow management, ease of configuration, and cross-platform support.

To fill these gaps, we present RSVrecon, a computational pipeline for assembling and analyzing sequencing data of RSV from clinical samples. The pipeline performs accurate genome reconstruction using optimized reference-guided assembly, followed by automated genotyping and phylogenetic analysis to characterize genetic diversity. RSVrecon includes specialized modules for identifying amino acid substitutions in the F protein, detecting mutations in monoclonal antibody sites that could potentially lead to antibody resistance, and supporting clinical decision-making. Furthermore, while complementing conventional text-based results from existing pipelines, RSVrecon delivers integrated graphical outputs specifically tailored for clinical practitioners. Finally, by utilizing Nextflow’s advanced workflow management, the pipeline delivers high-performance computing capabilities including parallel processing, automated logging, interruption recovery, and reproducibility across computing environments and infrastructures.

## Materials and methods

### Workflow design

RSVrecon pipeline consists of multiple sequential steps by following reference-based strategy designed for comprehensive RSV genomic analysis. Starting from raw sequencing data in the form of paired-end FASTQ files, each sample undergoes a standardized sequence assembly pipeline. This includes quality control and adapter trimming, followed by reference selection, read mapping, base calling, and final sequence assembly. To extract clinically relevant information such as viral genotype and F protein mutations, RSVrecon implements additional customized downstream analyses. These include gene annotation, genotype classification, polymorphism detection, screening for F protein substitutions, and phylogenetic analysis. The workflow concludes with a comprehensive reporting module that compiles results from all analytical steps into multiple formats, ensuring accessibility and usability for diverse audiences. These include a CSV report designed for bioinformaticians, a printer-friendly PDF summary, and an HTML report tailored specifically for clinical researchers and medical professionals (Figure 1-A).

**Figure 1.**
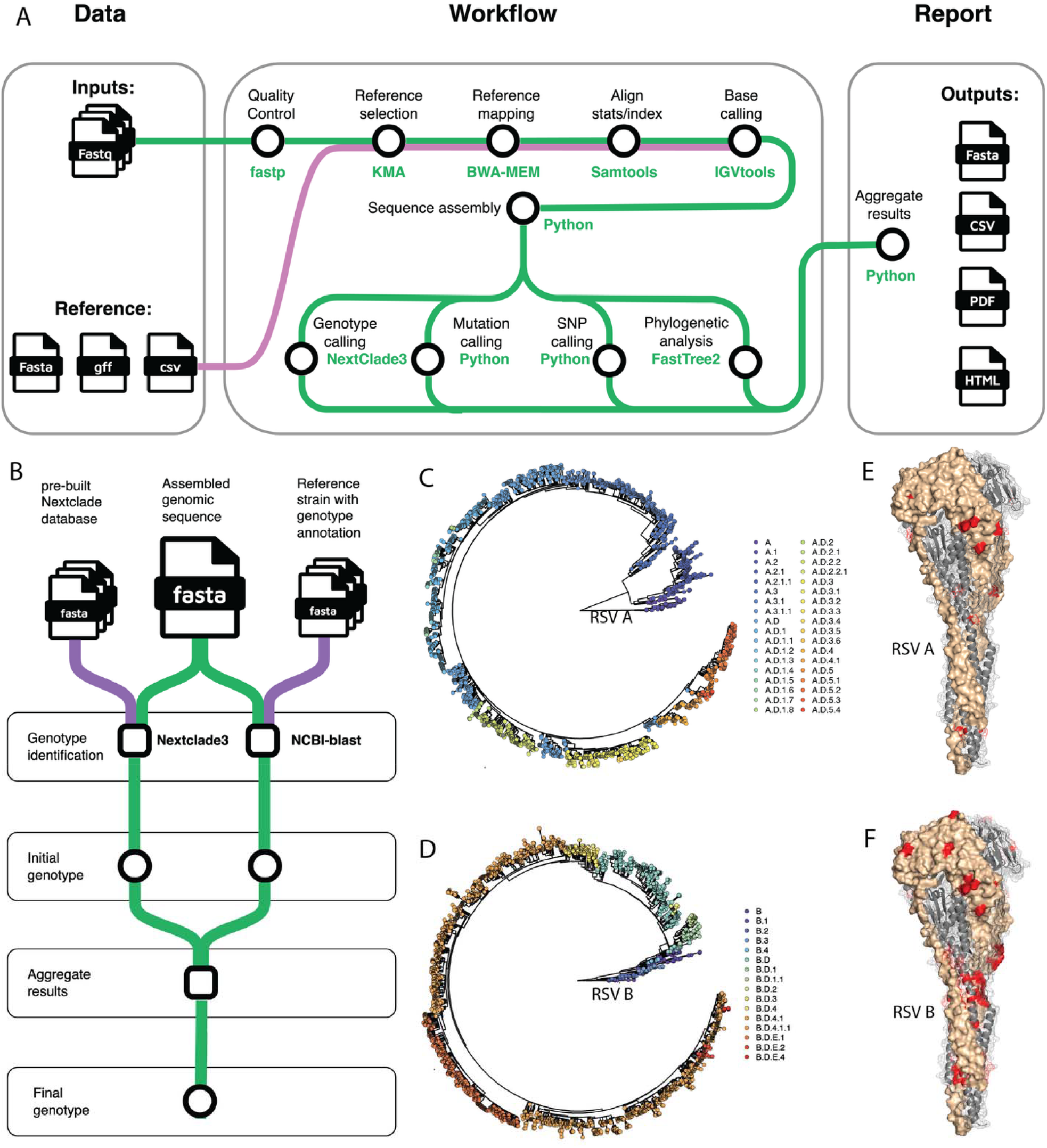
Workflow design and functionality of RSVrecon. **A**: Overview of the RSVrecon pipeline workflow. The pipeline takes raw sequence (paired FASTQ) as inputs from users. It then sequentially processes the data. Outputs include consensus sequences (FASTA) and aggregated results in multiple formats (CSV, PDF, HTML). The workflow is implemented in NextFlow, enabling parallelized execution and containerized reproducibility. Green paths represent the data flow of sequencing data through the pipeline, while purple paths indicate the data flow of pre-built reference datasets. **B**: workflow of genotype identification. Whole-genome-based genotypes are inferred using two approaches independently. Results from both approaches will be aggregated for a final call of genotype. **C, D**: Genotypes and genetic lineages of RSV A and RSV B defined using latest dataset. Phylogenetic trees are based on whole genome sequences. Trees and meta data are downloaded from Nextstrain (https://nextstrain.org/). **E, F**: Clinically relevant mutations in F protein for RSV A and RSV B, visualized on the structure of F protein in the post-fusion conformation (PDB ID: 3RRR).

Specifically, we utilized fastp (version 0.19.6) to trim raw paired read and remove adaptor sequences(Chen et al., 2018). The trimmed high-quality reads are then mapped against a comprehensive collection of RSV genomes by KMA (version 1.4.9) to select optimized reference genome for downstream analysis(Clausen et al., 2018). The optimized reference genome sequence are selected by balancing the highest ConClave score and longest reference coverage. After the reference genome being determined, the trimmed reads are precisely mapped against the reference genome by BWA (version 0.7.19-r127) (Li, 2013). The mapped results (SAM/BAM files) are handled by samtools (version 1.15.1) and the base counts are generated using IGVtools (version 2.3.2)(Li et al., 2009, Thorvaldsdóttir et al., 2013). Then the consensus sequence are generated from base count of each position, using a dual-coverage threshold of 10 and 50 (by default). While conventional tools typically use 10× coverage as default, we require ≥50× for high-confidence calls. For positions with 10-50× coverage, we apply an additional 90% nucleotide consensus requirement to ensure reliability, aiming to improve the detection of the dominant RSV population. Single-nucleotide polymorphisms (SNP) are calculated from base calling results, using a default cutoff of fraction 0.1. For each sample, phylogenetic analysis are performed using whole genome sequence to better reveal the viral evolution. Our pipeline selects two representative strains for each genetic lineages based on latest genotyping annotation and using their whole genome sequences as references in phylogenetics so that users can quickly know the evolutionary relationship between testing sample and existing lineages (Hadfield et al., 2018). We utilized FastTree2 to construct the maximum likelihood (ML) trees(Price et al., 2010). Lastly, we adopted NextClade3 and NCBI blast+ to infer genetic clade information(Aksamentov et al., 2021, McGinnis and Madden, 2004).

### Workflow implementation

To streamline installation and configuration for end users, we implemented this pipeline in Nextflow (https://www.nextflow.io/). Powered by Nextflow’s cross-platform property, our pipeline can be executed across different operating systems (Linux, macOS, Windows) and computing environments, including local computers, High Performance Computing (HPC) clusters, and cloud platforms (e.g., AWS). The pipeline leverages Nextflow’s DSL2 framework with modularized processes that maintain independent software environments, facilitating straightforward updates and maintenance. Dependency management is simplified through containerization technologies including Docker, Singularity, and Conda, eliminating the need for manual software installation and ensuring reproducible execution across diverse computational infrastructures. To enhance usability and foster community contributions, processes are integrated with the nf-core/modules repository whenever feasible. This implementation approach provides scalability and portability while maintaining the pipeline’s scientific reproducibility, allowing researchers to focus on RSV genomic analysis rather than technical configuration challenges.

For users that are unfamiliar with Nextflow, we also produced a Python-based implementation that utilizes Conda (https://conda.org/) for dependency management. We provided a bundled shell script that automates the environment setup and dependency installation to ensure the reproducibility across different environments and systems. The RSVrecon runtime environment is managed within an isolated Conda environment, which can be easily activated and deactivated, ensuring it remains independent from other system components. Once configured, the pipeline can be executed within this isolated environment in either local or distributed cluster mode. This is achieved through an integrated Python script that automatically manages sample queues and optimizes the use of available computational resources.

### Genotype Identification for Rapidly Evolving RSV

Identifying genotypes remains an ongoing effort for rapidly evolving viruses like RSV. An early study identified three (A1–A3) and seven (B1–B7) major genotypes for subtypes A and B, respectively, from RSV strains collected before February 2018 (Goya et al., 2020). A more recent study refined genotype classifications using data through March 2023, identifying a new major genotype group, A.D, for subtype A and reorganizing subtype B strains into five major groups: B.1–B.4, and B.D(Goya et al., 2024). We adopted NextClade (Version 3) as the primary method to identify and assign specific genotypes (genetic lineage level) for each tested sample (Aksamentov et al., 2021). It integrates a dynamically curated reference database sourced from the RSV Genotyping Consensus Consortium’s public repository (https://github.com/rsv-lineages). The database undergoes continuous updates to incorporate emerging genetic variants of RSV-A and RSV-B, reflecting current lineage definitions of 24 RSV-A and 16 RSV-B subtypes (Goya et al., 2024). It aligns the consensus sequence of the tested sample against the reference database, identifying all mutations in the query sequence relative to the reference sequences. Using these identified mutations, it searches for the best match on a pre-built phylogenetic tree and determines the appropriate genetic lineages to which the query sequence belongs. While this method has proven effective and efficient, it occasionally returns null results for certain query sequences, particularly for RSV samples from earlier years. To address such cases, we developed an alternative approach to assign a less precise genotype to the query sequence. The query sequence is mapped to a comprehensive, well-annotated collection of reference sequences using NCBI Blast (version 2.6.0), respectively(McGinnis and Madden, 2004). The genotype of the best-matching record is then assigned to the query sequence. Genotypes identified through this alternative method are marked with an asterisk (“*”) to notify users and prevent confusion.

### Comprehensive Annotation and Detection of Clinically Relevant F Protein Substitutions

Our pipeline annotates the coding sequences (CDS) of all 11 RSV genes for the assembled sequence of each sample based on the gene annotation file (GFF file) of the selected reference strain from GenBank(Sayers et al., 2022). This enables the extraction of the CDS region for each gene and the translation of nucleotide (NT) sequences into amino acid (AA) sequences. The tested samples are subsequently classified into two distinct groups: RSV A and RSV B. Samples demonstrating negative results, characterized by insufficient coverage across all genes, are excluded from further analysis. For each group, we collected key substitutions in the F protein that impact clinical treatment from literature (Table S1). Most of these are single-residue mutations, while a few involve co-occurring mutations at multiple residues (co-mutations). For single mutations, we screen each residue of the tested sequences to identify the presence of these key substitutions. For co-mutations, we scan combinations of residues involved in the co-mutation and identify their presence. This approach ensures comprehensive detection of clinically relevant substitutions in the F protein.

## Results

### Pipeline Architecture: Modular Design for Robust RSV Genomic Reconstruction

RSVrecon is designed for comprehensive genomic analysis of RSV from clinical samples, integrating quality control, genome assembly, variant calling, genotyping, and phylogenetic inference into a robust and reproducible workflow. Unlike conventional pipelines that terminate with sequence assembly, RSVrecon addresses critical gaps in RSV genomic surveillance by automating identification of clinically important features including genotypes and F-protein mutations. The pipeline provides an end-to-end solution that directly connects sequencing data generation with clinical utility, delivering actionable insights in formats accessible to end-users (Figure 1-A).

The pipeline accepts raw sequencing data (FASTQ) with flexible nomenclature to accommodate diverse study designs. It incorporates automated quality control and read trimming to ensure data integrity, followed by alignment (BWA-MEM), base calling (Samtools/IGVtools), and consensus generation using a reference-based mapping approach. Our pipeline implements a dual-coverage threshold system (10× and 50×) to optimize variant calling accuracy. Beyond assembly, RSVrecon performs genotyping, F-protein mutation screening, SNP calling, and phylogenetic analysis to extract clinically relevant information for end-users.

RSVrecon delivers high-quality consensus sequences alongside comprehensive analytical outputs including QC summaries, mapping statistics, phylogenetic relationships, and mutation profiles. Results are presented in both CSV format for further analysis and clinician-friendly graphical reports (PDF/HTML), featuring batch-level summaries and individual sample details. By integrating these analyses into a single workflow, the pipeline eliminates intermediate processing steps, significantly reducing turnaround time while maintaining rigorous standards to support time-sensitive clinical decisions (Figure 1-A).

### Screening of genotyping and F Protein Substitutions: Key Features for Clinical Decision-Making and Surveillance

We prioritize two clinically significant features essential for decision-making and surveillance: genotypes and F protein substitutions. Our pipeline determines whole-genome-based genotype for each sample. We adopted NextClade3 as the primary method to identify and assign specific genotypes (genetic lineage level) for each tested sample (Aksamentov et al., 2021). To handle cases where our primary method fails (particularly with older sequences), we implemented a secondary BLAST analysis that provides approximate genotype calls, clearly flagged with “*” to indicate reduced confidence (Figure 1-B). We then conduct phylogenetic analysis by constructing an approximately-maximum-likelihood tree that integrates the sample sequences with a reference set representing major genotypes and genetic lineages (Figure S1), enabling users to readily explore the evolutionary relationships between the tested samples and existing strains in the natural reservoir (Figure 1-C,D).

Key substitutions in the F protein can affect efficacy of mAb, and therefore were a primary focus of our pipeline. For example, residues V447 and K433 are located in the interaction region between the MK-1654 mAb and the RSV-F protein (Tang et al., 2019). Similarly, residues 62–69 and 196–212 in the RSV-F protein serve as binding sites for nirsevimab (formerly referred to as MEDI8897 in earlier literature) (Zhu et al., 2018, Zhu et al., 2017). Additionally, residues 254– 277 in the RSV-F protein are recognized as binding epitopes for palivizumab, while residues 161–182 are associated with suptavumab binding (Simões et al., 2021). Substitutions at these residues can potentially disrupt antibody binding, leading to resistance against the corresponding mAbs (Langedijk et al., 2022). Some mutations are subtype-specific; for instance, in RSV B, S190F, S211N, and S389P have become dominant since 2020(Holland et al., 2023, LaVerriere et al., 2025, Rios-Guzman et al., 2024, Yunker et al., 2024). Additionally, certain mutations may co-occur, further influencing viral evolution. For example, I64T not only has individual significance but also frequently co-occurs with K68E (Wilkins et al., 2023, Ahani et al., 2023). To ensure comprehensive, sensitive, and accurate screening, we maintain an up-to-date list of clinically significant F protein substitutions for both RSV subtypes, as reported in literature (Table S1). This list is fully customizable, allowing users to define specific co-mutation patterns for tailored and advanced analyses (Figure 1-E,F). Clinically important F protein substitutions are prominently highlighted in all report formats. Additionally, we extract amino acid variations at these key residues for all tested strains, enabling researchers to focus on variation patterns specific to these critical positions.

### Delivering Accessible and User-Friendly Outputs for Clinical Research Needs

We strive to provide highly accessible and interpretable results tailored to the needs of diverse researchers, especially clinical practice. While our pipeline delivers mapped results in BAM format and assembled sequences in FASTA format—both standard and widely accepted in the research community—it goes beyond these basics by offering user-friendly outputs designed for varied research purposes. In addition to standard outputs, RSVrecon highlights clinically critical genomic features and presents them in multiple formats: a summarized CSV report, an integrative PDF report, and an interactive HTML report (Figure 2). The CSV report includes essential details such as quality control metrics, reference mapping summaries, sequence quality, gene coverage, genotypes, and F protein mutations. However, graphical outputs like phylogenetic trees, coverage maps, and SNP visualizations cannot be incorporated into the CSV format. The PDF and HTML reports, on the other hand, provide comprehensive coverage of all results for each tested sample. The PDF report is optimized for printing, with well-structured tables and graphics, making it especially useful for clinicians and record-keeping. The HTML report offers an interactive experience, allowing computer users to navigate seamlessly through sample analyses and locate specific records with ease. Together, these features ensure our pipeline delivers both depth and accessibility, meeting the needs of researchers and clinicians alike.

**Figure 2.**
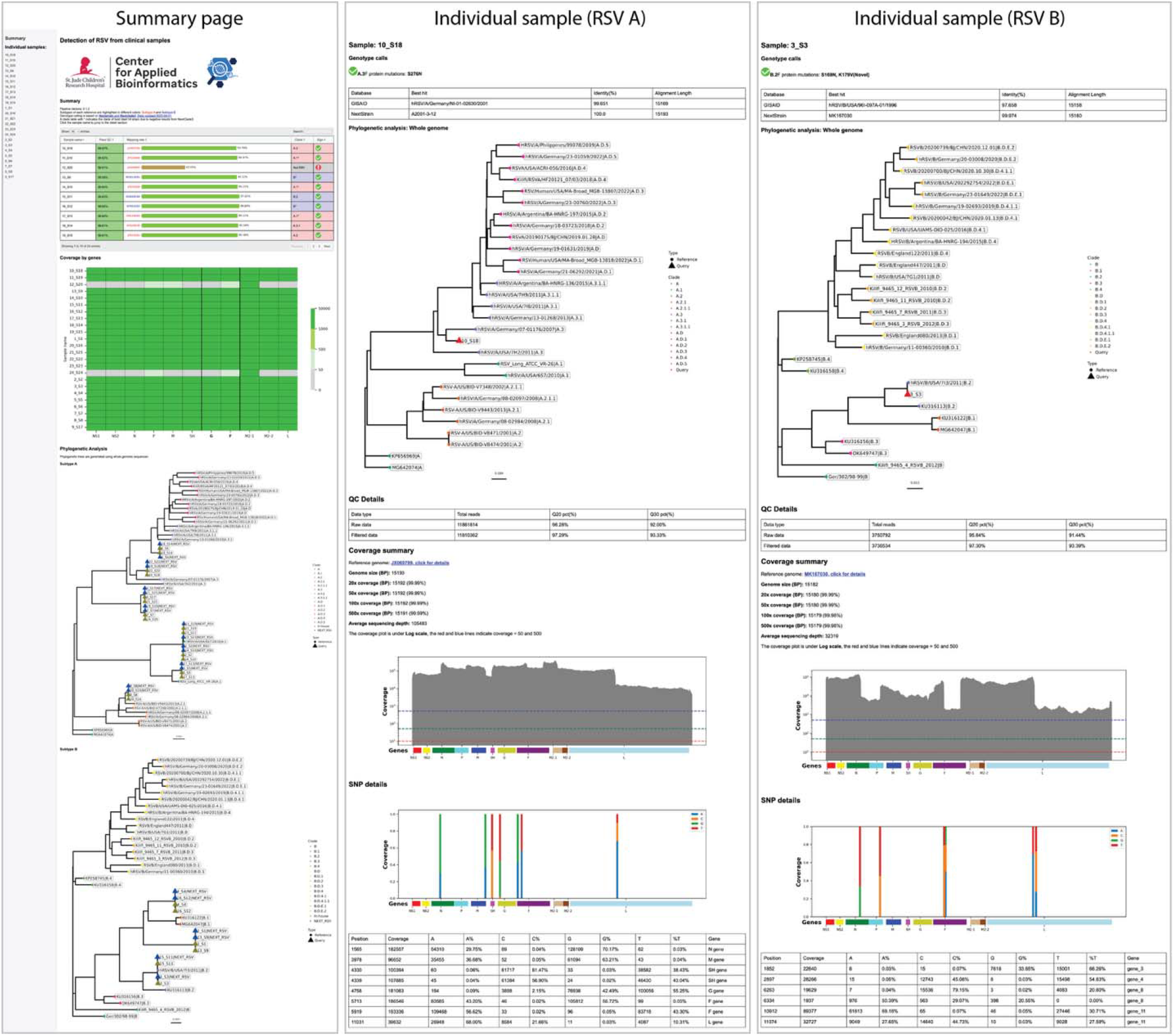
Accessible and User-Friendly Outputs of RSVrecon. The pipeline generates two levels of comprehensive reports: (1) a batch-level summary displaying QC metrics, mapping quality, subtypes, genotypes, gene coverage profiles, and phylogenetic relationships across all samples; and (2) detailed sample-specific reports featuring assembly quality metrics, genotype characterization, F protein mutations, best-matching reference sequences, whole-genome phylogenetic placement, coverage statistics, and SNP annotations.

### Scalable, robust and cross-platform system design and implementation

Beyond accuracy and functional capabilities, the technical implementation of bioinformatics pipelines critically determines their usability. RSVrecon employs Nextflow to ensure a robust, reproducible, and scalable bioinformatics workflow, combining standardized and custom analytical steps within a modular and maintainable pipeline structure. The pipeline follows nf-core’s (https://nf-co.re/) best practices for clear logic, organized file handling, and ease of debugging, re-using nf-core’s modules for upstream processes (e.g., data QC, read trimming, genome assembly) while incorporating custom downstream analyses (e.g., genotyping, SNP calling, F-protein mutation scanning, and phylogenetic analysis) for specialized clinical-relevant characterization for RSV (Ewels et al., 2020). To guarantee reproducibility across platforms, each step runs in an isolated containerized environment, leveraging Docker (https://www.docker.com/) for local PCs and Singularity (https://apptainer.org/) for HPC systems, ensuring robust performance and consistent results regardless of computational infrastructure. By merging Nextflow’s workflow management with containerization and a balanced approach to standardization and flexibility, RSVrecon delivers a high-quality, portable solution for RSV genomic research and surveillance. To ensure accessibility for users unfamiliar with Nextflow, we also developed a Python-based alternative that uses Conda (https://anaconda.org/) for reproducible dependency management, with a bundled script that automates setup and supports pipeline execution in both local and cluster environments.

### Comparison between RSVrecon and existing pipelines

We first benchmarked assembly accuracy, a fundamental aspect of genomic surveillance, between RSVrecon and existing RSV pipelines. Results indicated that all pipelines yielded comparable assembly accuracy for both subtypes (Figure S1-A,B). Then our comparative evaluation of RSVrecon and existing pipelines was structured around three key aspects: ***Functionality, Results Presentation***, and ***Usability***. Existing pipelines exhibit notable limitations in functionality: NEXT-RSV-PIPE focuses exclusively on sequence assembly without downstream analysis, while RSV-GenoScan offers F protein mutations scanning and restricted genotyping of G gene sequences, which has been deprecated recently (see supplementary for details)(Goya et al., 2024). In contrast, RSVrecon provides a comprehensive solution through (1) up-to-date genotyping using whole-genome sequences, (2) phylogenetic tree construction with reference strains for evolutionary insights, and (3) systematic detection of clinically relevant F protein mutations across all samples. Leveraging comprehensive analytical functions, researchers can uncover critical insights from RSV clinical samples with unprecedented clarity (Table 1).

**Table 1.**
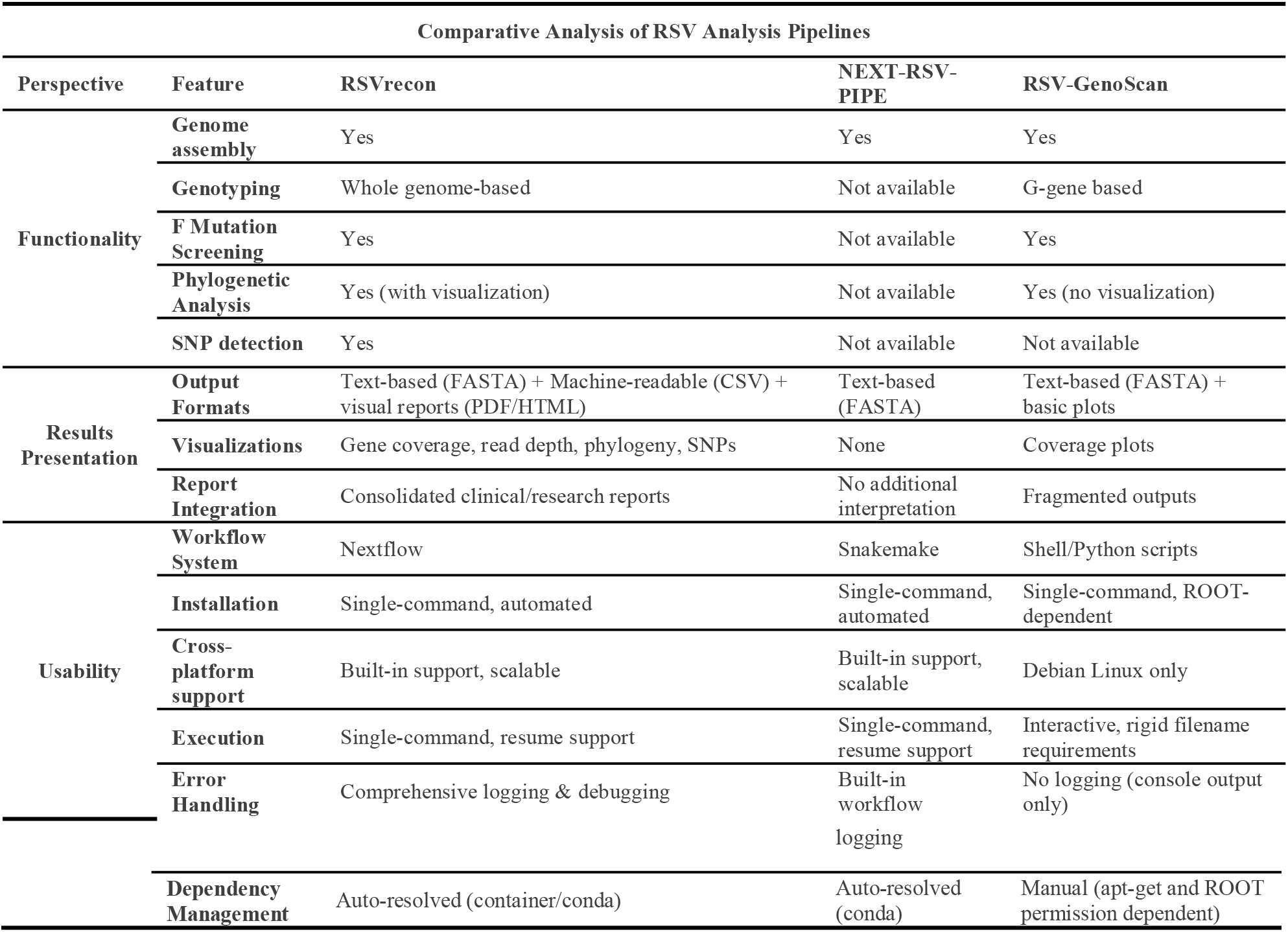
Comparison between RSVrecon and existing pipelines.

Regarding results presentation, existing pipelines primarily deliver non-integrated, text-based results (e.g., FASTA, GFF, or mutation lists), whereas RSVrecon enhances usability by combining standardized machine-readable outputs with intuitive visualizations. While NEXT-RSV-PIPE provides raw assembled sequences and annotations without interpretation, and RSV-GenoScan supplements its textual outputs with only basic coverage plots and a manual Newick-format phylogenetic tree, RSVrecon delivers both computational outputs and user-friendly visualizations. These include graphical representations of gene coverage, read depth, and phylogenetic relationships, alongside consolidated PDF/HTML reports tailored for clinical and research applications. By balancing bioinformatics rigor with end-user accessibility, RSVrecon bridges a critical gap in RSV data analysis and clinical application (Table 1).

In terms of usability, RSVrecon (Nextflow-based) and NEXT-RSV-PIPE (Snakemake-based) exemplify modern workflow design, featuring automated dependency management, environment compatibility, and robust error handling. Conversely, RSV-GenoScan’s reliance on root-dependent shell scripts and Debian-specific installations restricts its applicability, while its interactive execution requirements and inflexible filename conventions hinder scalability. RSVrecon and NEXT-RSV-PIPE support single-command execution and resume functionality— features absent in RSV-GenoScan. Additionally, RSVrecon and NEXT-RSV-PIPE leverage built-in logging and debugging, while RSV-GenoScan lacks logging, directing mixed messages to the console. These technical shortcomings significantly diminish RSV-GenoScan’s utility for large-scale NGS analysis compared to its workflow-managed counterparts (Table 1).

To enable systematic comparison, we quantified performance across 14 key features of three major categories and visualized the results as a radar plot (Figure 3; see Supplementary for scoring criteria). Our quantitative assessment demonstrates RSVrecon’s consistent superiority across all three evaluation dimensions, achieving maximum scores (5/5) in 12 of 14 features. While NEXT-RSV-PIPE shows limitations in functionality (missing 4/5 key analytical features) and results presentation (lacking visualizations), and RSV-GenoScan underperforms in usability (manual dependency management, OS restrictions) and report integration, RSVrecon unifies comprehensive analysis capabilities within a robust, user-friendly platform. The pipeline advances current standards by combining: (1) standardized computational outputs with publication-ready visualizations, overcoming the text-heavy outputs of existing tools; (2) a Nextflow-based architecture ensuring scalability, reproducibility, and automated error handling—addressing critical weaknesses of script-based alternatives like RSV-GenoScan; and (3) integrated clinical reporting that bridges the gap between computational analysis and biological interpretation. These innovations establish RSVrecon as the premier solution for modern RSV genomic surveillance and translational research.

**Figure 3.**
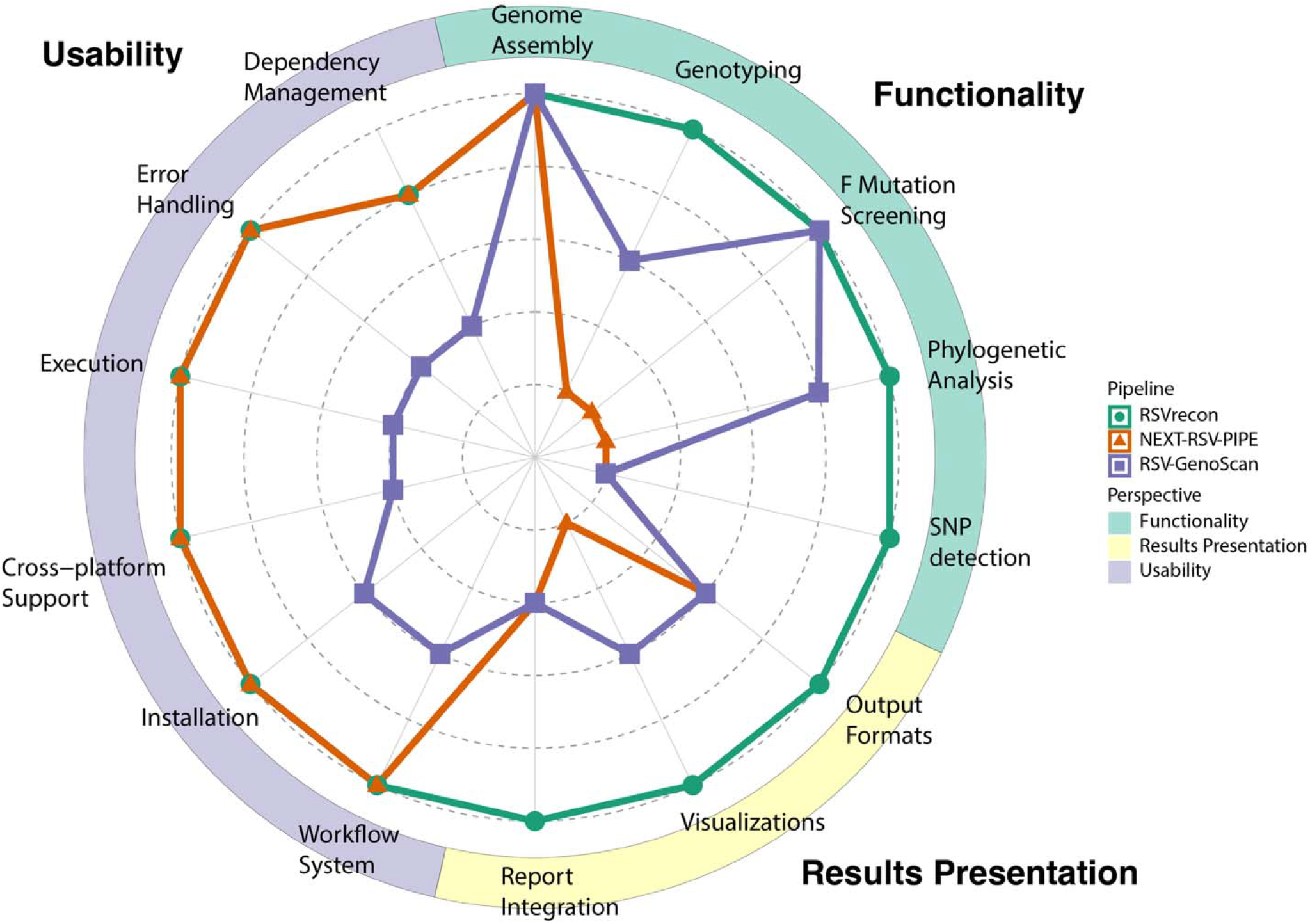
Comparative Analysis of RSV Analysis Pipelines. The radar plot evaluates three RSV analysis tools (RSVrecon, NEXT-RSV-PIPE, and RSV-GenoScan) across 14 features grouped into three perspectives: Functionality (light blue), Results Presentation (light yellow), and Usability (light purple). Each feature was scored on a 1-5 scale. RSVrecon demonstrates the most comprehensive capabilities, particularly in functionality and visualization, while NEXT-RSV-PIPE performs well in workflow management aspects. RSV-GenoScan shows strengths in specific functional components but has usability limitations.

## Discussion

Viral genome assembly from next-generation sequencing (NGS) data represents a well-established field, with mature computational workflows now achieving high accuracy in consensus generation. However, most existing pipelines—including those specifically optimized for RSV—remain narrowly focused on assembly, delivering raw genomic sequences (FASTA) as terminal outputs. While these results provide the essential foundation for downstream analyses, they are rarely actionable for end-users like clinicians and biologists, who require interpreted biological insights rather than technical sequence data. This persistent “last-mile gap” in data translation critically limits the real-world utility of genomic surveillance, particularly for time-sensitive RSV clinical management.

The clinical imperative is clear: RSV infection progression is rapid, and diagnostic delays directly compromise patient outcomes. Timely detection of key genomic features—including subtypes (RSV-A/B), F-protein antigenic variants, and known resistance markers—is essential to guide therapy, infection control, and public health responses. Conventional workflows, which decouple assembly from clinical interpretation, introduce bottlenecks that undermine this urgency. Our pipeline, RSVrecon, addresses this limitation by unifying genome reconstruction with downstream functional annotation, delivering clinician-friendly reports (PDF/HTML) that highlight actionable features such as genotype-phenotype correlations and vaccine-escape mutations. By collapsing the traditional multi-step analysis cascade into a single automated workflow, we reduce turnaround time and eliminate dependency on bioinformatic intermediaries, ensuring genomic insights directly inform bedside decisions.

Researchers have diverse priorities and requirements for presenting sequencing data, depending on their focus. Bioinformaticians and biostatisticians, who often perform large-scale analyses using computational tools, prefer standardized, machine-readable formats such as FASTA for assembled sequences, BAM for mapping results, and CSV/TSV for metadata. In contrast, clinicians and biologists, who manually review cases to guide clinical decisions, value user-friendly report formats like PDFs for easy printing and record-keeping, HTML reports for computer-based reviews, and CSV files for integrating clinical and research records. However, existing pipelines primarily cater to computational needs, overlooking the practical requirements of clinicians, making it challenging for them to access essential information efficiently. Our pipeline addresses this issue by providing results in diverse formats, ensuring accessibility and usability for all users, with a particular emphasis on meeting clinical needs.

In conclusion, our pipeline offers a robust framework for RSV genomic analysis, seamlessly integrating sequential steps to deliver high-quality results and clinically actionable insights. From processing raw sequencing data to producing comprehensive reports, each stage is carefully designed to maximize accuracy and ease of use. Key features include precise genome assembly, reliable genotype determination, and customizable F protein substitution analysis, meeting essential clinical and research needs. By integrating advanced tools, utilizing a curated GenBank reference repository, and maintaining a comprehensive catalog of key F protein mutations, the pipeline ensures accurate, sensitive, and up-to-date genotype assignments and mutation analysis. The final reporting function synthesizes these findings into user-friendly formats suitable for bioinformaticians, clinical researchers, and medical professionals alike. Furthermore, the modular design supports future development, enabling applications to other viruses and expanding its utility. This combination of accuracy, flexibility, and user-focused design establishes our pipeline as a valuable resource for decision-making, surveillance, and advancing RSV research.

Notably, all three RSV-specific pipelines evaluated in this study employ reference-mapping-based approaches. This design choice is driven by the clinical need to derive consensus sequences representing the dominant viral strain in a sample, which is critical for diagnostic decision-making. However, co-infections—where multiple viral genotypes coexist—are common in clinical practice. For epidemiological and infectious disease research, quantifying the relative abundances of these genotypes provides valuable insights. While reference mapping reliably captures the dominant variant, it may overlook minority strains. In contrast, de novo assembly can reconstruct multiple contigs, revealing the genetic diversity arising from co-infections (Zerbino and Birney, 2008). Additionally, tools like kallisto (https://github.com/pachterlab/kallisto) (Bray et al., 2016) enable lineage abundance quantification by transcript-level profiling, an approach successfully applied to SARS-CoV-2 wastewater surveillance (Baaijens et al., 2022). Integrating such methods into RSV pipelines could address current limitations, offering a more comprehensive analysis of clinical samples by characterizing both dominant and minor viral populations.

## Supporting information

supplemental meterial

## Acknowledgement

This research included experiments conducted by the Center for Applied Bioinformatics (CAB) which is supported in part by ALSAC and the National Cancer Institute grant P30 CA021765.

## Code availability

Nextflow pipeline is available at https://github.com/stjudecab/rsvrecon and the python pipeline is available at https://github.com/stjudecab/RSVreconPy.

## Author contribution

Conceptualization: L.L., H.Y., and G.W.; Methodology: L.L.; Investigation: L.L., G.W., D.R.H.; Supervision: L.L.; Implementation: L.L., H.Y.; Experiment: J.N.B., R.T.H., D.R.H.; Writing – original draft: L.L., H.Y.; Writing – review & editing: L.L., H.Y., J.N.B., R.W., R.T.H., G.W., and D.R.H.

## References

Ahani, B., Tuffy, K. M., Aksyuk, A. A., Wilkins, D., Abram, M. E., Dagan, R., Domachowske, J. B., Guest, J. D., Ji, H. & Kushnir, A. 2023. Molecular and phenotypic characteristics of RSV infections in infants during two nirsevimab randomized clinical trials. Nature Communications, 14, 4347.

Aksamentov, I., Roemer, C., Hodcroft, E. B. & Neher, R. A. 2021. Nextclade: clade assignment, mutation calling and quality control for viral genomes. Journal of open source software, 6, 3773.

Antipov, D., Rayko, M., Kolmogorov, M. & Pevzner, P. A. 2022. viralFlye: assembling viruses and identifying their hosts from long-read metagenomics data. Genome biology, 23, 57.

Baaijens, J. A., Zulli, A., Ott, I. M., Nika, I., Van Der Lugt, M. J., Petrone, M. E., Alpert, T., Fauver, J. R., Kalinich, C. C. & Vogels, C. B. 2022. Lineage abundance estimation for SARS-CoV-2 in wastewater using transcriptome quantification techniques. Genome biology, 23, 236.

Battles, M. B. & Mclellan, J. S. 2019. Respiratory syncytial virus entry and how to block it. Nature Reviews Microbiology, 17, 233–245.

Biggerstaff, M., Cauchemez, S., Reed, C., Gambhir, M. & Finelli, L. 2014. Estimates of the reproduction number for seasonal, pandemic, and zoonotic influenza: a systematic review of the literature. BMC infectious diseases, 14, 1–20.

Bray, N. L., Pimentel, H., Melsted, P. & Pachter, L. 2016. Near-optimal probabilistic RNA-seq quantification. Nature biotechnology, 34, 525–527.

Chen, S., Zhou, Y., Chen, Y. & Gu, J. 2018. fastp: an ultra-fast all-in-one FASTQ preprocessor. Bioinformatics, 34, i884–i890.

Chevreux, B. 2007. MIRA: an automated genome and EST assembler.

Clausen, P. T., Aarestrup, F. M. & Lund, O. 2018. Rapid and precise alignment of raw reads against redundant databases with KMA. BMC bioinformatics, 19, 1–8.

Dawood, F. S., Iuliano, A. D., Reed, C., Meltzer, M. I., Shay, D. K., Cheng, P.-Y., Bandaranayake, D., Breiman, R. F., Brooks, W. A. & Buchy, P. 2012. Estimated global mortality associated with the first 12 months of 2009 pandemic influenza A H1N1 virus circulation: a modelling study. The Lancet infectious diseases, 12, 687–695.

Dosbaa, A., Guilbaud, R., Yusti, A.-M. F., Ferré, V. M., Charpentier, C., Descamps, D., LE Hingrat, Q. & CoppéE, R. 2024. RSV-GenoScan: An automated pipeline for whole-genome human respiratory syncytial virus (RSV) sequence analysis. Journal of Virological Methods, 327, 114938.

Ewels, P. A., Peltzer, A., Fillinger, S., Patel, H., Alneberg, J., Wilm, A., Garcia, M. U., DI Tommaso, P. & Nahnsen, S. 2020. The nf-core framework for community-curated bioinformatics pipelines. Nature biotechnology, 38, 276–278.

Falsey, A. R., Hennessey, P. A., Formica, M. A., Cox, C. & Walsh, E. E. 2005. Respiratory syncytial virus infection in elderly and high-risk adults. New England Journal of Medicine, 352, 1749–1759.

Fu, P., Wu, Y., Zhang, Z., Qiu, Y., Wang, Y. & Peng, Y. 2024. VIGA: a one-stop tool for eukaryotic virus identification and genome assembly from next-generation-sequencing data. Briefings in Bioinformatics, 25, bbad444.

Goldstein, S. A., Brown, J., Pedersen, B. S., Quinlan, A. R. & Elde, N. C. 2022. Extensive recombination-driven coronavirus diversification expands the pool of potential pandemic pathogens. Genome Biology and Evolution, 14, evac161.

Goya, S., Galiano, M., Nauwelaers, I., Trento, A., Openshaw, P. J., Mistchenko, A. S., Zambon, M. & Viegas, M. 2020. Toward unified molecular surveillance of RSV: A proposal for genotype definition. Influenza and Other Respiratory Viruses, 14, 274–285.

Goya, S., Ruis, C., Neher, R. A., Meijer, A., Aziz, A., Hinrichs, A. S., Von Gottberg, A., Roemer, C., Amoako, D. G. & Acuña, D. 2024. Standardized phylogenetic classification of human respiratory syncytial virus below the subgroup level. Emerging Infectious Diseases, 30, 1631.

Hadfield, J., Megill, C., Bell, S. M., Huddleston, J., Potter, B., Callender, C., Sagulenko, P., Bedford, T. & Neher, R. A. 2018. Nextstrain: real-time tracking of pathogen evolution. Bioinformatics, 34, 4121–4123.

Han, L., Li, L., Wen, F., Zhong, L., Zhang, T. & Wan, X.-F. 2019. Graph-guided multi-task sparse learning model: a method for identifying antigenic variants of influenza A (H3N2) virus. Bioinformatics, 35, 77–87.

Holland, L. A., Holland, S. C., Smith, M. F., Leonard, V. R., Murugan, V., Nordstrom, L., Mulrow, M., Salgado, R., White, M. & Lim, E. S. 2023. Genomic sequencing surveillance to identify respiratory syncytial virus mutations, Arizona, USA. Emerging Infectious Diseases, 29, 2380.

Jansz, N. & Faulkner, G. J. 2024. Viral genome sequencing methods: benefits and pitfalls of current approaches. Biochemical Society Transactions, BST20231322.

Kaler, J., Hussain, A., Patel, K., Hernandez, T. & Ray, S. 2023. Respiratory syncytial virus: a comprehensive review of transmission, pathophysiology, and manifestation. Cureus, 15.

Köndgen, S., Oh, D.-Y., Thürmer, A., Sedaghatjoo, S., Patrono, L. V., Calvignac-Spencer, S., Biere, B., Wolff, T., Dürrwald, R. & Fuchs, S. 2024. A robust, scalable, and cost-efficient approach to whole genome sequencing of RSV directly from clinical samples. Journal of Clinical Microbiology, 62, e01111–23.

Langedijk, A. C., Harding, E. R., Konya, B., Vrancken, B., Lebbink, R. J., Evers, A., Willemsen, J., Lemey, P. & Bont, L. J. 2022. A systematic review on global RSV genetic data: Identification of knowledge gaps. Reviews in medical virology, 32, e2284.

Laverriere, E., Behar, S., Sher-Jan, C., Liang, Y. M., Sagar, M. & Connor, J. H. Genomic Epidemiology of Respiratory Syncytial Virus in a New England Hospital System, 2024. Open Forum Infectious Diseases, 2025. Oxford University Press US, ofaf334.

Li, H. 2013. Aligning sequence reads, clone sequences and assembly contigs with BWA-MEM. arXiv preprint 1303.3997.

Li, H., Handsaker, B., Wysoker, A., Fennell, T., Ruan, J., Homer, N., Marth, G., Abecasis, G., Durbin, R. & Subgroup, G. P. D. P. 2009. The sequence alignment/map format and SAMtools. bioinformatics, 25, 2078–2079.

Li, L., Chang, D., Han, L., Zhang, X., Zaia, J. & Wan, X.-F. 2020a. Multi-task learning sparse group lasso: a method for quantifying antigenicity of influenza A (H1N1) virus using mutations and variations in glycosylation of Hemagglutinin. BMC bioinformatics, 21, 1–22.

Li, X., Giorgi, E. E., Marichannegowda, M. H., Foley, B., Xiao, C., Kong, X.-P., Chen, Y., Gnanakaran, S., Korber, B. & Gao, F. 2020b. Emergence of SARS-CoV-2 through recombination and strong purifying selection. Science advances, 6, eabb9153.

Mcginnis, S. & Madden, T. L. 2004. BLAST: at the core of a powerful and diverse set of sequence analysis tools. Nucleic acids research, 32, W20–W25.

Morens, D. M. & Fauci, A. S. 2007. The 1918 influenza pandemic: insights for the 21st century. The Journal of infectious diseases, 195, 1018–1028.

Neal, H. E., Barrett, C. T., Edmonds, K., Moncman, C. L. & Dutch, R. E. 2024. Examination of respiratory syncytial virus fusion protein proteolytic processing and roles of the P27 domain. Journal of Virology, 98, e01639–24.

Petrova, V. N. & Russell, C. A. 2018. The evolution of seasonal influenza viruses. Nature Reviews Microbiology, 16, 47–60.

Price, M. N., Dehal, P. S. & Arkin, A. P. 2010. FastTree 2–approximately maximum-likelihood trees for large alignments. PloS one, 5, e9490.

QIAGEN 2025. CLC Genomics Workbench, QIAGEN Digital Insights.

Rios-Guzman, E., Simons, L. M., Dean, T. J., Agnes, F., Pawlowski, A., Alisoltanidehkordi, A., Nam, H. H., Ison, M. G., Ozer, E. A. & Lorenzo-Redondo, R. 2024. Deviations in RSV epidemiological patterns and population structures in the United States following the COVID-19 pandemic. Nature communications, 15, 3374.

Sayers, E. W., Cavanaugh, M., Clark, K., Pruitt, K. D., Schoch, C. L., Sherry, S. T. & Karsch-Mizrachi, I. 2022. GenBank. Nucleic acids research, 50, D161–D164.

Simões, E. A., Forleo-Neto, E., Geba, G. P., Kamal, M., Yang, F., Cicirello, H., Houghton, M. R., Rideman, R., Zhao, Q. & Benvin, S. L. 2021. Suptavumab for the prevention of medically attended respiratory syncytial virus infection in preterm infants. Clinical Infectious Diseases, 73, e4400–e4408.

Smith, D. J., Lapedes, A. S., De Jong, J. C., Bestebroer, T. M., Rimmelzwaan, G. F., Osterhaus, A. D. & Fouchier, R. A. 2004. Mapping the antigenic and genetic evolution of influenza virus. science, 305, 371–376.

Tang, A., Chen, Z., Cox, K. S., Su, H.-P., Callahan, C., Fridman, A., Zhang, L., Patel, S. B., Cejas, P. J. & Swoyer, R. 2019. A potent broadly neutralizing human RSV antibody targets conserved site IV of the fusion glycoprotein. Nature communications, 10, 4153.

Thorvaldsdóttir, H., Robinson, J. T. & Mesirov, J. P. 2013. Integrative Genomics Viewer (IGV): high-performance genomics data visualization and exploration. Briefings in bioinformatics, 14, 178–192.

Váradi, A., Kaszab, E., Kardos, G., Prépost, E., Szarka, K. & Laczkó, L. 2022. Rapid genotyping of targeted viral samples using Illumina short-read sequencing data. Plos one, 17, e0274414.

Wan, Y., Renner, D. W., Albert, I. & Szpara, M. L. 2015. VirAmp: a galaxy-based viral genome assembly pipeline. Gigascience, 4, s13742-015-0060-y.

Wang, H., Paulson, K. R., Pease, S. A., Watson, S., Comfort, H., Zheng, P., Aravkin, A. Y., Bisignano, C., Barber, R. M. & Alam, T. 2022. Estimating excess mortality due to the COVID-19 pandemic: a systematic analysis of COVID-19-related mortality, 2020–21. The Lancet, 399, 1513–1536.

Wilkins, D., Langedijk, A. C., Lebbink, R. J., Morehouse, C., Abram, M. E., Ahani, B., Aksyuk, A. A., Baraldi, E., Brady, T. & Chen, A. T. 2023. Nirsevimab binding-site conservation in respiratory syncytial virus fusion glycoprotein worldwide between 1956 and 2021: an analysis of observational study sequencing data. The Lancet Infectious Diseases, 23, 856–866.

Yamashita, A., Sekizuka, T. & Kuroda, M. 2016. VirusTAP: viral genome-targeted assembly pipeline. Frontiers in microbiology, 7, 32.

Yang, X., Charlebois, P., Gnerre, S., Coole, M. G., Lennon, N. J., Levin, J. Z., Qu, J., Ryan, E. M., Zody, M. C. & Henn, M. R. 2012. De novo assembly of highly diverse viral populations. BMC genomics, 13, 1–13.

Yunker, M., Fall, A., Norton, J. M., Abdullah, O., Villafuerte, D. A., Pekosz, A., Klein, E. & Mostafa, H. H. 2024. Genomic Evolution and Surveillance of Respiratory Syncytial Virus during the 2023–2024 Season. Viruses, 16, 1122.

Zerbino, D. R. & Birney, E. 2008. Velvet: algorithms for de novo short read assembly using de Bruijn graphs. Genome research, 18, 821–829.

Zhu, Q., Lu, B., Mctamney, P., Palaszynski, S., Diallo, S., Ren, K., Ulbrandt, N. D., Kallewaard, N., Wang, W. & Fernandes, F. 2018. Prevalence and significance of substitutions in the fusion protein of respiratory syncytial virus resulting in neutralization escape from antibody MEDI8897. The Journal of infectious diseases, 218, 572–580.

Zhu, Q., Mclellan, J. S., Kallewaard, N. L., Ulbrandt, N. D., Palaszynski, S., Zhang, J., Moldt, B., Khan, A., Svabek, C. & Mcauliffe, J. M. 2017. A highly potent extended half-life antibody as a potential RSV vaccine surrogate for all infants. Science translational medicine, 9, eaaj1928.

Zhu, Q., Patel, N. K., Mcauliffe, J. M., Zhu, W., Wachter, L., Mccarthy, M. P. & Suzich, J. A. 2012. Natural polymorphisms and resistance-associated mutations in the fusion protein of respiratory syncytial virus (RSV): effects on RSV susceptibility to palivizumab. Journal of Infectious Diseases, 205, 635–638.

Zsichla, L., Zeeb, M., Fazekas, D., Áy, É., Müller, D., Metzner, K. J., Kouyos, R. D. & Müller, V. 2024. Comparative Evaluation of Open-Source Bioinformatics Pipelines for Full-Length Viral Genome Assembly. Viruses, 16, 1824.

