## supplemental meterial for "Bridging Genomics and Clinical Medicine: RSVrecon Enhances RSV Surveillance with Automated Genotyping and Clinically Important Mutation Reporting"

### Supplementary

**Dual-coverage cutoff ensures accuracy of genome assembly**

The cost of next-generation sequencing (NGS) has significantly decreased over the past decades, making it standard practice to improve accuracy by increasing sequencing depth. For example, in the early NGS era (2008–2015), a few million reads per sample were typically sufficient for bulk RNA-seq analysis ([McGettigan, 2013](https://pmc.ncbi.nlm.nih.gov/articles/PMC2949280/); [Wang et al., 2009](https://pmc.ncbi.nlm.nih.gov/articles/PMC3031867/)). However, current best practices often require 100–200 million reads per sample to ensure precise gene expression quantification and facilitate complex analyses such as alternative splicing detection. For viral sequencing, particularly for RNA viruses with short but highly mutable genomes—such as Influenza viruses and respiratory syncytial virus (RSV)—a high sequencing coverage is essential for accurate genome assembly and variant detection. A minimum average coverage of 100× is generally required, with over 1000× being common to capture low-frequency variants and maintain accuracy ([Hoffmann et al., 2022](https://www.mdpi.com/2076-393X/10/8/1359); [Formenti et al., 2023](https://onlinelibrary.wiley.com/doi/10.1111/irv.13252)). As sequencing depth continues to increase, it is crucial to adjust coverage cutoffs accordingly to maintain data reliability and prevent erroneous results.

To assess the impact of coverage thresholds, we applied our pipeline alongside the RSV-specific pipelines to a batch of sequencing data. The results demonstrated that both pipelines produced consistent outcomes for most samples in both subtypes (Figure S1-A,B). However, a low default coverage cutoff can occasionally yield inaccurate results for samples with in-sufficient number of reads. RSVrecon implemented a dual-coverage threshold system to optimize variant calling accuracy. The lower cutoff is 10x and the higher cutoff is 50x by default (adjustable). Comparing to a single cutoff system, it balances sensitivity and accuracy of the assembly, ensure the quality of results.

While RSVrecon (Nextflow) and RSV-NEXT-PIPE (Snakemake) support dynamic parameterization—enabling users to define coverage cutoffs via configuration files—RSV-GenoScan relies on hardcoded values, restricting optimization for specific use cases. While users have the option to modify these parameters, in practice, most tend to rely on the default settings. Therefore, establishing well-calibrated default parameters is crucial. We advocate for a conservative approach to default parameterization, as overly aggressive settings may lead to erroneous results, particularly in cases involving poor-quality samples, without providing adequate warnings. A conservative default configuration or dual-coverage threshold system ensures accurate results while triggering errors in specific scenarios, prompting users to adjust parameters appropriately.

**Scoring criteria for pipeline evaluation**

We developed a quantitative scoring system (1-5 scale, each scale represents: 5: Comprehensive implementation with advanced capabilities, 4: Functional implementation with minor limitations, 3: Partial/basic implementation, 2: Minimal functionality, 1: Feature absent) to evaluate pipelines across 14 features grouped into three categories:

**A. Functionality**

1. Genome Assembly

- 5: Automated, optimized assembly
- 1: No assembly capability

2. Genotyping

- 5: Whole-genome based with updated reference database
- 3: G-gene only with limited references
- 1: Not available

3. F Mutation Screening

- 5: Automated detection with clinical annotation
- 1: No screening capability

4. Phylogenetic Analysis

- 5: Automated tree construction with visualization
- 4: Tree construction without visualization
- 1: Not available

5. SNP Detection

- 5: Genome-wide variant calling
- 1: No SNP detection

**B. Results Presentation**

6. Output Formats
   - 5: Machine-readable (CSV/FASTA) + visual reports (PDF/HTML)
   - 3: Text-based only (FASTA/GFF)
   - 1: Non-standardized outputs
7. Visualizations
   - 5: Interactive multi-plot outputs
   - 3: Static basic plots
   - 1: No visualizations
8. Report Integration
   - 5: Consolidated clinical/research reports
   - 2: Fragmented outputs

**C. Usability**

9. Technical Implementations

- 5: Fully automated best practices (e.g., containerized, resume support)
- 3: Partial automation
- 1: Manual configuration required

**Genotype reference of RSV**

We utilized the latest genotype annotations from the RSV Genotyping Consensus Consortium’s public repository (<https://github.com/rsv-lineages>)(Goya et al., 2024). This resource provides curated genotype designations for both RSV subtypes A and B.

Between January 2024 and March 2025, the Consortium released four updates for Subtype A (2024-01-16, 2024-01-29, 2024-08-01, 2024-11-27) and five for Subtype B (2024-01-16, 2024-01-29, 2024-08-01, 2024-11-27, 2025-03-04). Notably, the 2025-03-04 release for Subtype B formally deprecated and removed the G-gene genotype designation.

RSVrecon sources these annotations from the NextClade data repository (https://github.com/nextstrain/nextclade_data), which synchronizes with the Consortium’s releases, enabling genotyping via NextClade3.

**Publicly available resources of RSV**

Several publicly available resources provide valuable data on respiratory syncytial virus (RSV), forming the foundation of this study. GenBank (https://www.ncbi.nlm.nih.gov/genbank/) offers a comprehensive collection of genomic sequences across various species, including RSV. The Global Initiative on Sharing All Influenza Data (GISAID) (https://gisaid.org/), originally established for avian influenza data, has expanded to include other respiratory viruses such as COVID-19, dengue, influenza, Mpox, and RSV, providing not only genomic sequences but also insights into RSV’s evolutionary dynamics, epidemiological surveillance, and statistical trends. Nextstrain (https://nextstrain.org/) tracks the evolution of various pathogens, including RSV, by aggregating strains from GenBank and continuously updating phylogenetic and genotyping analyses. Additionally, The Protein Data Bank (PDB) (https://www.rcsb.org/) archives three-dimensional protein structures, and in this study, we retrieved the RSV F protein structure (PDB ID: 3RRR) to visualize clinically relevant mutations. These resources collectively enhance our understanding of RSV genomics, evolution, and structural biology, facilitating more robust analyses in this study.

**Data visualization**

Phylogenetic trees and meta data in Figure 1 were obtained from Nextstrain (<https://nextstrain.org/>), data updated 2024-07-12. Trees in Figure S1 and S2 were generated from genomic sequences. Sequences were aligned using MAFFT (version 7.490), and the approximately maximum-likelihood trees were generated using FastTree2 (version 2.1). Trees are visualized using ggtree (version 3.6.2), ggplot2 (version 3.4.2) and treeio (version 1.22.0) in R (version 4.2.2). Structure of F protein is obtained from PDB, PDB ID is 3RRR. The structure was visualized using PyMol (version 2.6).

**References**

Wang, Zhong, Mark Gerstein, and Michael Snyder. "RNA-Seq: a revolutionary tool for transcriptomics." *Nature reviews genetics* 10, no. 1 (2009): 57-63.

Ozsolak, Fatih, and Patrice M. Milos. "RNA sequencing: advances, challenges and opportunities." *Nature reviews genetics* 12, no. 2 (2011): 87-98.

Galli, Cristina, Erika Ebranati, Laura Pellegrinelli, Martina Airoldi, Carla Veo, Carla Della Ventura, Arlinda Seiti et al. "From clinical specimen to whole genome sequencing of a (H3N2) influenza viruses: a fast and reliable high-throughput protocol." *Vaccines* 10, no. 8 (2022): 1359.

Wang, Xinye, Ki Wook Kim, Gregory Walker, Sacha Stelzer‐Braid, Matthew Scotch, and William D. Rawlinson. "Genome characterization of influenza A and B viruses in New South Wales, Australia, in 2019: A retrospective study using high‐throughput whole genome sequencing." *Influenza and Other Respiratory Viruses* 18, no. 1 (2024): e13252.

GOYA, S., RUIS, C., NEHER, R. A., MEIJER, A., AZIZ, A., HINRICHS, A. S., VON GOTTBERG, A., ROEMER, C., AMOAKO, D. G. & ACUÑA, D. 2024. Standardized phylogenetic classification of human respiratory syncytial virus below the subgroup level. Emerging Infectious Diseases, 30, 1631.

**Table S1**. Clinically relevant mutations identified from literatures.

|  | **Mutations** | **Clinical impact** | **Source** |
| --- | --- | --- | --- |
| **Subtype A** | E66K | nirsevimab | Langedijk et al., 2022; Zhu et al., 2017; Zhu et al., 2018 |
|  | I206T | nirsevimab | Langedijk et al., 2022; Zhu et al., 2017; Zhu et al., 2018 |
|  | S255N | palivizumab | Langedijk et al., 2022; Simões et al., 2021 |
|  | D263N | palivizumab | Langedijk et al., 2022; Simões et al., 2021 |
|  | K272M\|E | palivizumab | Langedijk et al., 2022; Simões et al., 2021 |
|  | S276K\|N\|R | palivizumab | Langedijk et al., 2022; Simões et al., 2021 |
|  | V447M | MK‐1654 | Langedijk et al., 2022; Tang et al., 2019 |
|  | N165K | suptavumab | Langedijk et al., 2022; Simões et al., 2021 |
|  | V178L | suptavumab | Langedijk et al., 2022; Simões et al., 2021 |
| **Subtype B** | R42K | palivizumab | Zhu et al., 2012 |
|  | I64T | nirsevimab | Langedijk et al., 2022; Zhu et al., 2017; Zhu et al., 2018 |
|  | K65E | nirsevimab | Langedijk et al., 2022; Zhu et al., 2017; Zhu et al., 2018 |
|  | T67N | nirsevimab | Langedijk et al., 2022; Zhu et al., 2017; Zhu et al., 2018 |
|  | K68N\|R\|Q\|E | nirsevimab | Langedijk et al., 2022; Zhu et al., 2017; Zhu et al., 2018 |
|  | S169N | suptavumab | Langedijk et al., 2022; Simões et al., 2021 |
|  | L172Q | suptavumab | Langedijk et al., 2022; Simões et al., 2021 |
|  | S173L | suptavumab | Langedijk et al., 2022; Simões et al., 2021 |
|  | K176R | suptavumab | Langedijk et al., 2022; Simões et al., 2021 |
|  | K179I | suptavumab | Langedijk et al., 2022; Simões et al., 2021 |
|  | S190N |  | Holland et al., 2023; LaVerriere et al., 2025; Rios-Guzman et al., 2024 |
|  | N200D | nirsevimab | Langedijk et al., 2022; Zhu et al., 2017; Zhu et al., 2018 |
|  | N\|K201S\|K | nirsevimab | Langedijk et al., 2022; Zhu et al., 2017; Zhu et al., 2018 |
|  | L204S | nirsevimab | Langedijk et al., 2022; Zhu et al., 2017; Zhu et al., 2018 |
|  | I206M | nirsevimab | Langedijk et al., 2022; Zhu et al., 2017; Zhu et al., 2018 |
|  | N208S\|D | nirsevimab | Langedijk et al., 2022; Zhu et al., 2017; Zhu et al., 2018 |
|  | Q209K\|L\|R | nirsevimab | Langedijk et al., 2022; Zhu et al., 2017; Zhu et al., 2018 |
|  | S211N | nirsevimab | Langedijk et al., 2022; Zhu et al., 2017; Zhu et al., 2018 |
|  | S255G | palivizumab | Langedijk et al., 2022; Simões et al., 2021 |
|  | M264I | palivizumab | Langedijk et al., 2022; Simões et al., 2021 |
|  | K272N\|Q\|R | palivizumab | Langedijk et al., 2022; Simões et al., 2021 |
|  | L273I | palivizumab | Langedijk et al., 2022; Simões et al., 2021 |
|  | S276N | palivizumab | Langedijk et al., 2022; Simões et al., 2021 |
|  | S389P | palivizumab | Holland et al., 2023; LaVerriere et al., 2025; Rios-Guzman et al., 2024; Yunker et al., 2024 |
|  | K433R | MK‐1654 | Langedijk et al., 2022; Tang et al., 2019 |
|  | I64T+K68E | nirsevimab | Wilkins et al., 2023 |

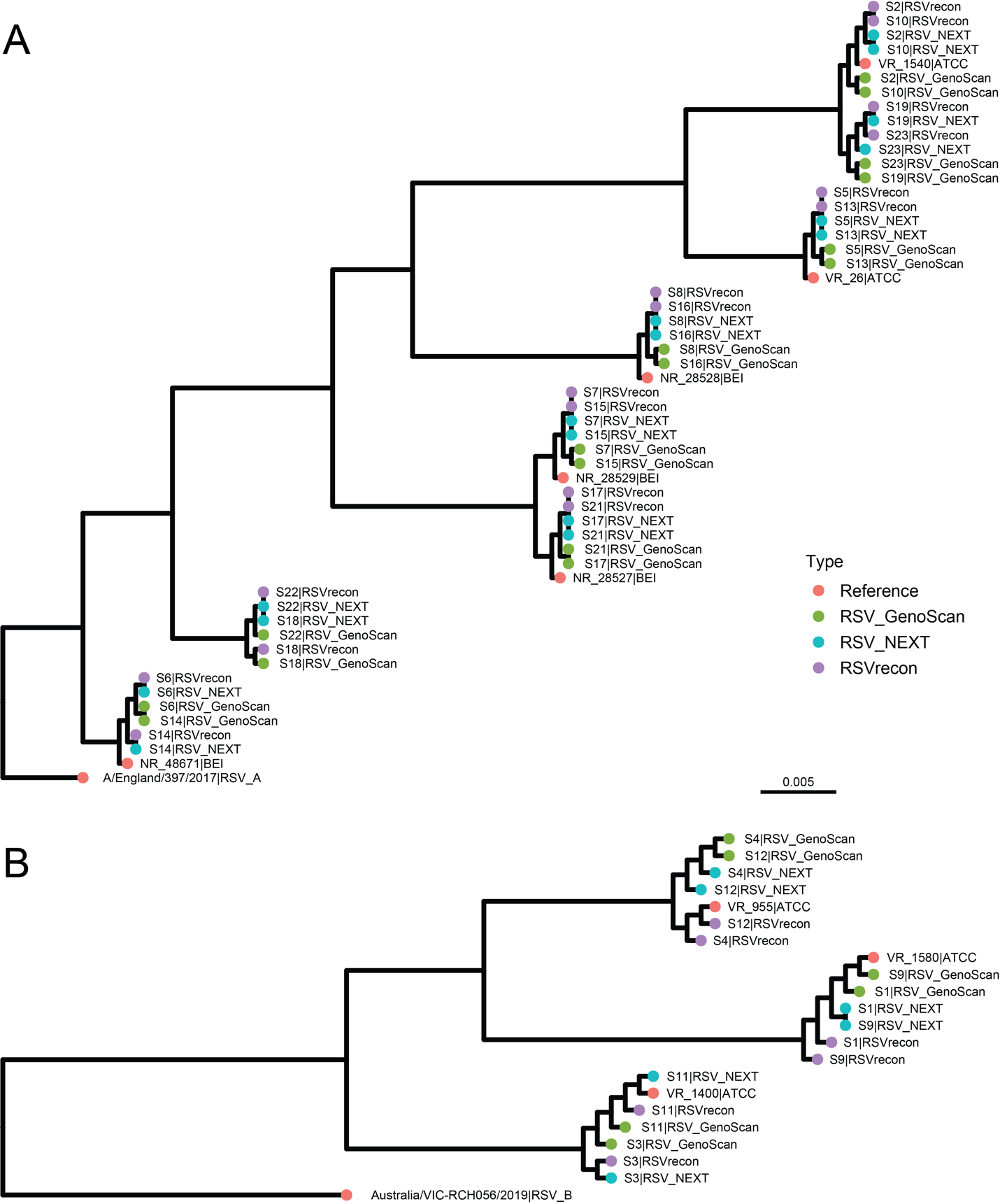

**Figure S1**. Phylogenetic tree including assembled sequences and reference. Sequences generated from different pipelines are indicated by colors. A: Phylogenetic tree of tested samples of subtype A. B: Phylogenetic tree of tested samples of subtype B.
